# PyHFO: Lightweight Deep Learning-powered End-to-End High-Frequency Oscillations Analysis Application

**DOI:** 10.1101/2023.08.25.554741

**Authors:** Yipeng Zhang, Lawrence Liu, Yuanyi Ding, Xin Chen, Tonmoy Monsoor, Atsuro Daida, Shingo Oana, Shaun Hussain, Raman Sankar, Fallah Aria, Jerome Engel, Richard J. Staba, William Speier, Jianguo Zhang, Hiroki Nariai, Vwani Roychowdhury

## Abstract

In the context of epilepsy studies, intracranially-recorded interictal high-frequency oscillations (HFOs) in EEG signals are emerging as promising spatial neurophysiological biomarkers for epileptogenic zones. While significant efforts have been made in identifying and understanding these biomarkers, deep learning is carving novel avenues for biomarker detection and analysis. Yet, transitioning such methodologies to clinical environments is difficult due to the rigorous computational needs of processing EEG data via deep learning. This paper presents our development of an advanced end to end software platform, PyHFO, aimed at bridging this gap. PyHFO provides an integrated and user-friendly platform that includes time-efficient HFO detection algorithms such as short-term energy (STE) and Montreal Neurological Institute and Hospital (MNI) detectors and deep learning models for artifact and HFO with spike classification. This application functions seamlessly on conventional computer hardware. Our platform has been validated to adeptly handle datasets from 10-minute EEG recordings captured via grid/strip electrodes in 19 patients. Through implementation optimization, PyHFO achieves speeds up to 50 times faster than the standard HFO detection method. Users can either employ our pre-trained deep learning model for their analyses or use their EEG data to train their model. As such, PyHFO holds great promise for facilitating the use of advanced EEG data analysis tools in clinical practice and large-scale research collaborations.

## I. Introduction

**H**UMAN and animal studies of epilepsy have suggested that intracranially-recorded interictal high-frequency oscillations (HFOs) in EEG signals are a promising spatial neurophysiological biomarker of the epileptogenic zone. Many retrospective studies [1]–[4] demonstrated that the removal of brain regions producing HFOs correlated with post-operative seizure freedom. More recently, various studies [5]–[11] have suggested that HFOs potentially have different mechanistic origins, and hence, only a subset of HFO events –often referred to as pathological HFOs – constitute meaningful biomarkers for epileptic zones, while others of physiological origins might be useful for characterizing, for example, the eloquent cortices [11]. Such further refinements of HFOs include tasks, such as artifact rejection, HFOs with spike-wave discharges (spk-HFO) detection, epileptogenic HFO discovery, and physiological HFO detection.

However, translating these research findings into a clinical setting to enhance seizure-free surgery outcomes poses significant challenges. It requires a multidisciplinary approach involving experts in machine learning, artificial intelligence, neurology, and epileptology to refine and establish the clinical relevance of different types of HFOs and potentially discover more effective biomarkers. Such collaboration necessitates the development of a scalable software platform that enables advanced data analysis, annotation, expert verification, and sharing of patient outcome data in a user-friendly manner.

In the field of EEG studies, a considerable number of open-source software applications aim to offer visualization tools, such as EEGLab [12], EEGnet [13], EPViz [14], and Brainstorm [15], to facilitate researchers in visualizing EEG signals. Additionally, there is a substantial array of computational biomarker detection algorithms implemented in various programming languages such as MNE [16], YASA [17], and PyEEG [18], which equip users with technical backgrounds to detect and visualize biomarkers using software. More recently, significant efforts have been devoted to deep learning for event classification in both scalp EEG [19] and invasive EEG [20] facilitating EEG decoding [21], artifact rejection [22], and disease detection [23]. However, as the complexity of network architectures continues to increase, coupled with the burgeoning development within the deep-learning community, the computational cost has dramatically risen. Consequently, a considerable scientific and engineering chasm has emerged between the research on deep learning-powered EEG analysis tools and the distribution of state-of-the-art deep learning methods to clinicians’ personal computers for practical application. So far, efforts to bridge this gap have been insufficient.

A similar scenario prevails in HFO studies. RIPPLELAB [24], an open-source Matlab-based software, has facilitated early studies on HFOs, incorporating EEG visualization and mainstream HFO detection algorithms. This software is widely used in several studies across the community [25]–[30]. Simultaneously, many recent studies are endeavoring to leverage deep learning models to carry out HFO analysis [10], [11], [31]. The research community values open-source HFO-analysis software like RIPPLELAB; however, the absence of an integrated platform for clinicians to employ these deep learning models hinders the full clinical potential of HFOs and associated biomarkers.

Therefore, a software platform compatible with popular deep learning frameworks is highly desirable for enabling advanced machine learning and deep learning tools to automate various steps of HFO refinement and deploy them efficiently even on moderately powerful machines commonly available to clinicians.

In this paper, we present our initial efforts in developing such an application, addressing three key engineering challenges:

- We developed time-efficient detection algorithms of HFO events, by re-implementing the HFO detectors in Python and significantly reduced the detection run-time by at least 50 times compared to state-of-the-art with comparative study to ensure the correctness of our implementation compared to Matlab-based counterpart.
- We addressed the demanding task of integration of deep learning-based HFO classification by simplifying artifact and spk-HFO classification networks introduced in a previous study [10], allowing the deep learning model to run smoothly on the CPU of researchers’ personal machine
- We build an open-source executable software that integrates both time-efficient HFO detection algorithms and simplified artifact and spk-HFO classification networks,

The integration of all of the functionalities, PyHFO, holds great potential for facilitating seamless collaboration and enabling large-scale EEG data analysis.

## II. Method

PyHFO is a multi-window Graphical User Interface (GUI) desktop application specifically designed for the efficient analysis and classification of HFOs. It presents a user-friendly and intuitive interface that caters to both technical and non-technical users, streamlining the process of HFO detection and classification. PyHFO operates through four primary stages upon loading an EEG recording: EEG signal reading, data filtering, HFO detection, and Deep Learning (DL)-based HFO classification. These stages are detailed in a data flowchart, as seen in Figure 1. The output of this pipeline includes detected events based on the implemented detection algorithm, accompanied by annotations of real HFOs, artifacts, and HFO-with-spikes (spk-HFO), generated using pre-trained DL-based HFO classification models. The specifics of each critical stage are elaborated upon in the subsequent sections.

**Fig. 1.**
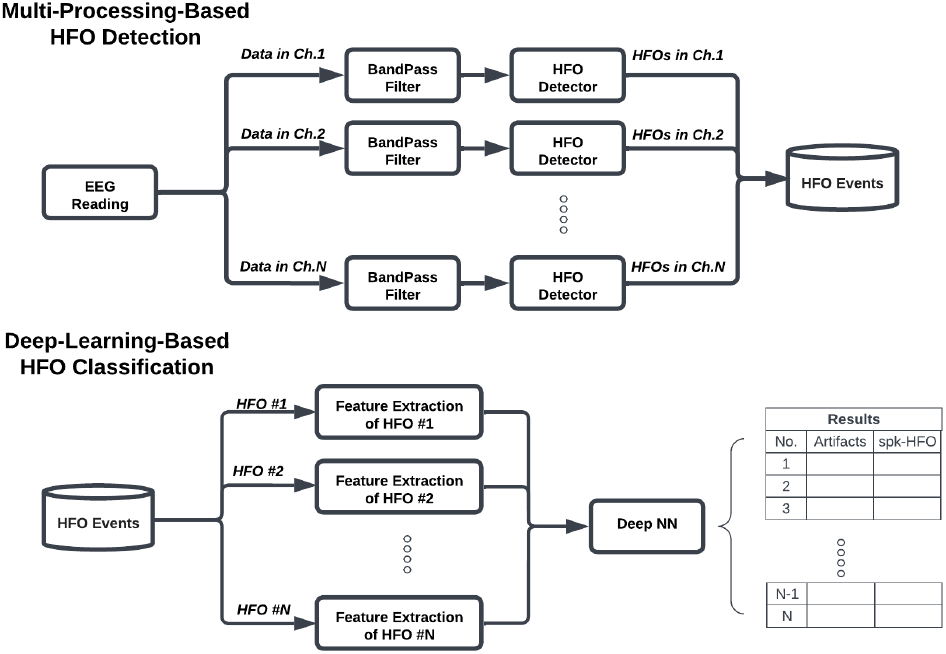
Our study’s overall data processing workflow is shown as a flowchart. We adopt a multi-processing mechanism in both HFO detection and feature extraction for deep learning networks, which significantly increases the efficiency of the HFO analysis.

### A. HFO detection algorithms

In PyHFO, we have integrated two automatic HFO detection algorithms, namely the Short Time Energy (STE) [32] and the Montreal Neurological Institute and Hospital detector (MNI) [33]. They are selected because they are the two primary detection algorithms in the widely used Matlab-based HFO analysis tool, RIPPLELAB, where they have demonstrated success in numerous studies. Thanks to PyHFO’s modular architecture, coupled with the open-source ethos of the project, allows for easy integration of alternative HFO detection methods if required. Developers simply need to adhere to a straightforward interface. Moreover, any added methods will inherently benefit from the established multi-processing paradigm.

We have faithfully replicated the precise parameters and computational implementation of both algorithms from RIPPLELAB in Python. The flowcharts of these two algorithms are in Figure 2 and Figure 3; we have put a detailed explanation in Appendix. However, we have introduced certain modifications. This includes replacing functions, such as the gamma distribution parameter estimation, with the official Scipy’s APIs. Additionally, we have exposed the random seed to the user to ensure the reproducibility of the MNI detector. More importantly, to enhance execution efficiency, we have replaced the for loop with matrix multiplication, particularly in the Gabor wavelet computation. These alterations may result in slightly divergent detection outcomes compared to those obtained from RIPPLELAB. A comprehensive analysis and comparison of these results will be presented in the dedicated analysis section.

**Fig. 2.**
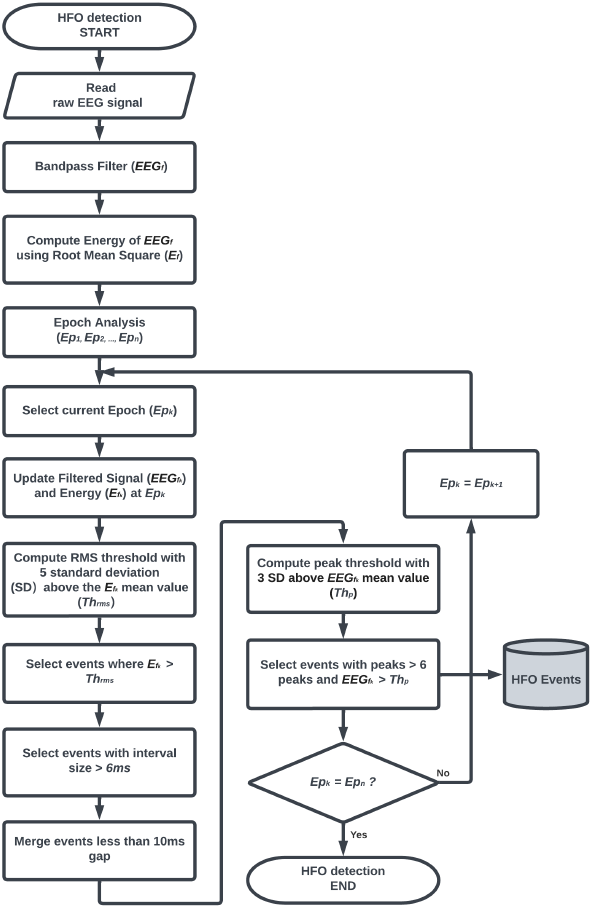
The computational flowchart of the STE HFO detection algorithm.

**Fig. 3.**
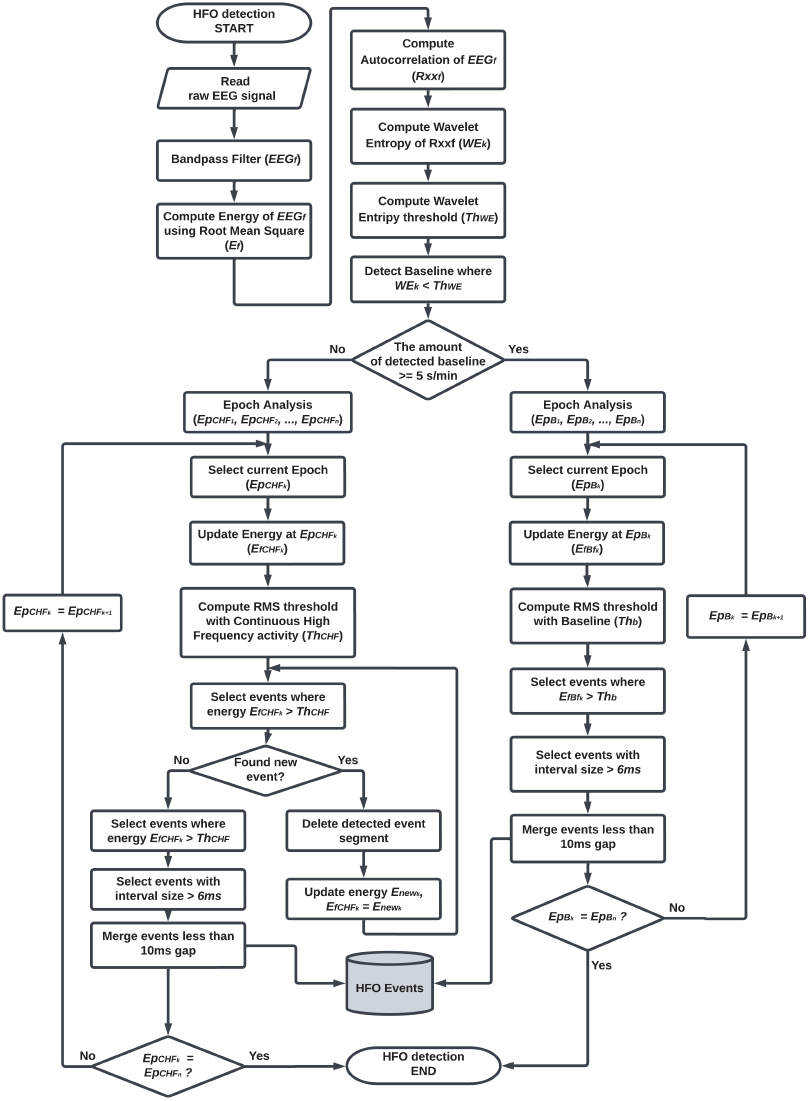
The computational flowchart of the MNI HFO detection algorithm.

### B. HFO detector implementation details

#### 1) Data reading

PyHFO is designed to accept mainstream EEG data file formats such as the European Data Format (EDF). Additionally, it can process data in the widely-used NumPy format when users employ the deployed Python package (see Section II-E). In processing an EDF file, raw data is stored in binary format. Upon reading, to convert the digital (raw) values *D*_*raw*_ to the real-world physical voltage *V*, the digital values are calibrated using maximum and minimum physical voltage *V*_*max*_, *V*_*min*_ and maximum and minimum digital values *D*_*max*_, *D*_*min*_ The equation used by most EEG data processing tools, such as MNE [16], is given in Equation (1). In this equation, *R* is the calibration ratio, defined as the ratio of the difference between the maximum and minimum physical values to the difference between the maximum and minimum digital values, and an offset *O* is defined as the difference between the minimum physical value and the product of the calibration ratio and the minimum digital value.

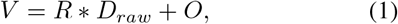

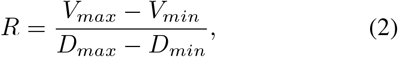

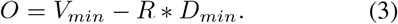

It’s worth noting that RIPPLELAB processes EDF files differently than other mainstream EDF reading tools. Contrastingly, RIPPLELAB performs calibration only through *V* = *R *D*_*raw*_, with no offset adjustment, resulting in data readings with a DC offset between RIPPLELAB and other EDF reading tools. In our implementation within PyHFO, we have elected to use the more robust calibration equation, Equation (1), as the open-sourced Python package MNE.

#### 2) Signal filtering

The voltage value read from EDF is then passed through a bandpass filter to extract the signal in the desired frequency domain with the specified ripple and attenuation. The bandpass filter used in the RIPPLELAB is the Chebyshev type II filter; the parameter of this filter consists of PassBand(*F*_*p*_), StopBand(*F*_*s*_), PassBand Ripple(*r*_*p*_), and StopBand Attenuation(*r*_*s*_). For constructing such a filter, the order of the filter is first estimated, and then the frequency response is constructed. We noticed that Matlab sometimes could not achieve an exact match to the desired PassBand Ripple and StopBand Attenuation. Therefore, in implementing the PyHFO, we choose to use the filter construction by Scipy as it can produce a more aligned frequency response to the specification. However, since the Scipy cannot replicate the filter parameter specified in RIPPLELAB, *F*_*p*_ = 80 Hz, *F*_*s*_ = 500 Hz, *r*_*p*_ = 0.5 dB, and *r*_*s*_ = 100 dB due to the numerical overflow. We used the closest number in StopBand Attenuation *r*_*s*_ = 93 dB instead in our implementation. We visualize such prementioned phenomena in Figure 4.

**Fig. 4.**
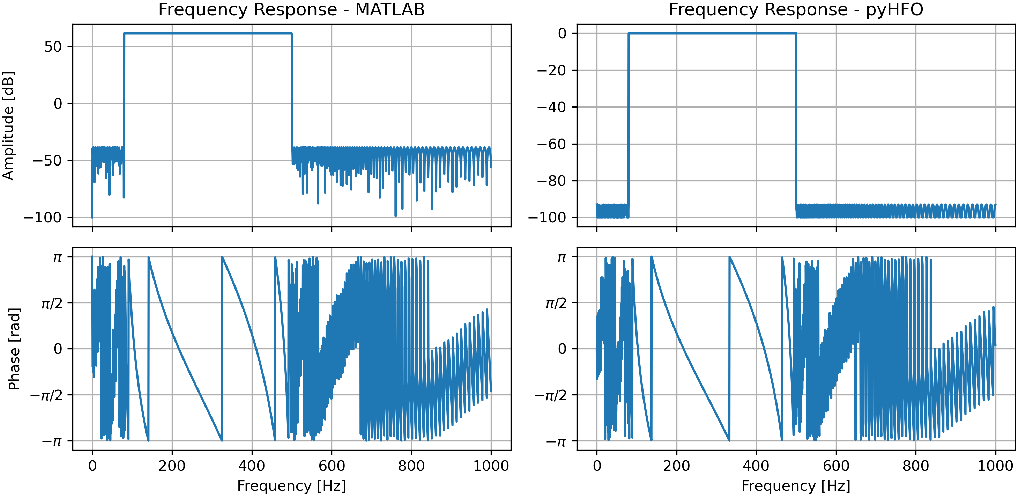
A comparison between Matlab and Scipy regarding filter construction. For the given parameters *F*_*p*_ = 80 Hz, *F*_*s*_ = 500 Hz, *r*_*p*_ = 0.5 dB, and *r*_*s*_ = 100 dB, which is specified in RIPPLELAB, Matlab did not precisely match the frequency response to the desired PassBand Ripple and StopBand Attenuation. In contrast, Scipy utilized in our implementation generates a more closely aligned frequency response. Owing to numerical overflow, we use a slightly adjusted StopBand Attenuation value of *r*_*s*_ = 93 dB in our PyHFO implementation.

#### 3) Multi-processing-based detection framework

To improve the efficiency of our HFO detection pipeline, we leverage the multi-processing capability of Python. In Figure 1, we illustrate how we parallelize the time-consuming steps, namely data filtering and HFO detection, across each channel. Since the computation for each channel is independent, we assign each channel’s data to different CPU processors to run simultaneously. This approach maximizes the use of available CPU cores, leading to a significant reduction in the detection time. To demonstrate the acceleration of our detector’s running speed, we compare the detection times for MNI and STE detectors using both our detector and the RIPPLELAB detector on different hardware machines.

### C. Lightweight deep learning-based HFO classification

We follow the same artifact and spk-HFO classifier design in [10] as it has already shown promising performance against expert annotation. The training data is from HFO detected by STE detector in 10 min along with annotation from experts [10]. However, several limitations are preventing them from being directly used in the natural setting: 1) Currently, the HFO analysis is majorly conducted in CPU machines; access to GPU is not very popular in this domain of study. Application of the proposed network in [10] in CPU is time-consuming. 2) Then the generalization ability of models in [10] cannot be ensured in HFOs detected by other detectors such as MNI. To address 1), we reduce the computational cost of the model by first seeking the smallest information (input size) that can keep the classification performance and then by employing the state-of-the-art neural network pruning technique to reduce the network size. To address limitation 2), we develop a data-augmentation strategy in the neural network training to improve the generalization ability of the model.

#### 1) DL model training with data augmentation

The time-domain augmentation is used to improve the generalization ability of the artifact and spk-HFO detector. During the training of the network, we randomly flip the EEG signals and randomly shift the center of the HFO event forward and backward 50 ms, as shown in Figure 5. We train and test the model performance on the annotated 10 min EEG data in [10] and reported 5-fold cross-validation results with a corresponding 95% confidence interval. Additionally, to ensure the DL models trained from the STE detector also generalize well in HFO events detected by the MNI detector, an expert (HN and SH) annotated MNI HFOs into the artifact, spk-HFO, non-spk-HFO from representative patients and the performance metric of the model against the expert annotation is reported. The inter-rater reliability of these two expert annotators is measured by the Cohen kappa score (kappa = 0.96 for labeling artifacts, 0.85 for labeling HFOs with spikes). The procedure of the evaluation was reported in [10].

**Fig. 5.**
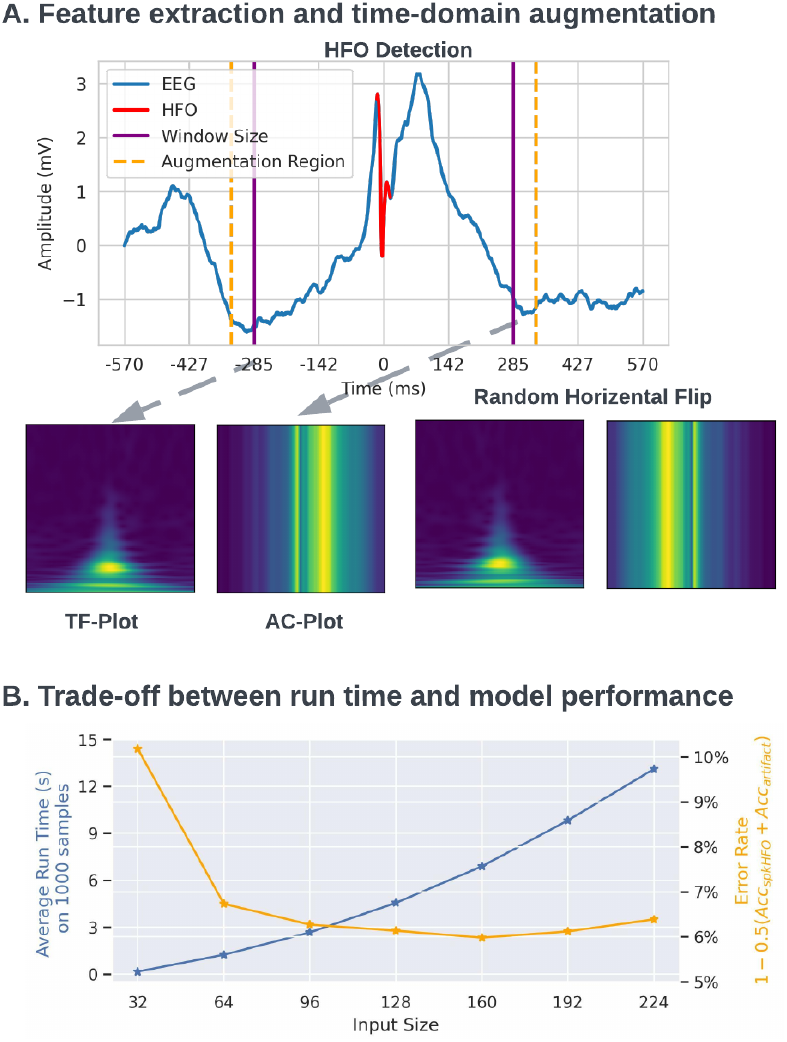
Deep learning network architecture and complexity; A. The time-domain augmentation consists of two steps, 1. center of the HFO is first randomly shifted by 50ms, before and after. 2, the EEG signal is randomly flipped in the time domain. Then the time-frequency plot (10 290 Hz) and amplitude coding plot with size 128x128 is generated. B. Empirical analysis of the model input size, network performance, and model complexity. While keeping the information resolution (2.18Hz/pix and 4.46ms/pix) the same, we train and evaluate the two classifiers with different input sizes from 32 x 32 (10 ∼ 80Hz, *±* 72 ms) to 224 x 224 (10 ∼ 500 Hz, *±*500 ms), we plot the error rate of these two classifiers and the average run time with the corresponding input size together. The error rate of these two classifiers is defined as 1 − 0.5 (Acc_spk-HFO_ + Acc_artifacts_) in 5-fold cross-validation and the average run time is the time to predict 1000 samples on CPU using a Linux Machine 10 times,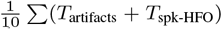. We chose 128 as the input size because it is the best tradeoff between speed and performance.

#### 2) Reduction of the computational cost

The computational cost of a neural network can be measured by the total number of Multiply Accumulate (MACs) for a fixed number of inputs, which is influenced by the dimension of the input and the size of the architecture. To reduce the computational complexity introduced in the input dimension, we first reduce the redundancy in the dimension of the input by only using one time-frequency plot for the artifact detector and concatenation of only the time-frequency plot and amplitude coding plot as input for the spk-HFO classifier. Then we reduced the input dimension from 224 * 224 to 128 * 128, by only taking 10 to 290 Hz in the frequency domain and *±*285 ms of the center of the event in the time domain; these values are chosen by empirical analysis of balancing the computational complexity of the neural network and the classification accuracy Figure 5. Then we simplified the architecture of the artifact and spk-HFO detector, respectively, by pruning the neural network using DeepGraph [34] as shown in Figure 6. The interactive pruning was conducted for 5000 iterations, and the model was fine-tuned every 250 iterations by five epochs. Finally, we imposed a rule-based filter by treating all HFOs detected in the beginning one second and last one second as artifacts because the beginning and the ending of the recording will lead to artifacts production. The simplified network should run at the best tradeoff between speed and performance in CPU, and we also enable the use of GPU for users with GPU access on their machine. We evaluate the model complexity by computing MACs using one data sample. Additionally, we measure the average running speed of the model inference on 1000 data samples to get a more straightforward overview of the model complexity.

**Fig. 6.**
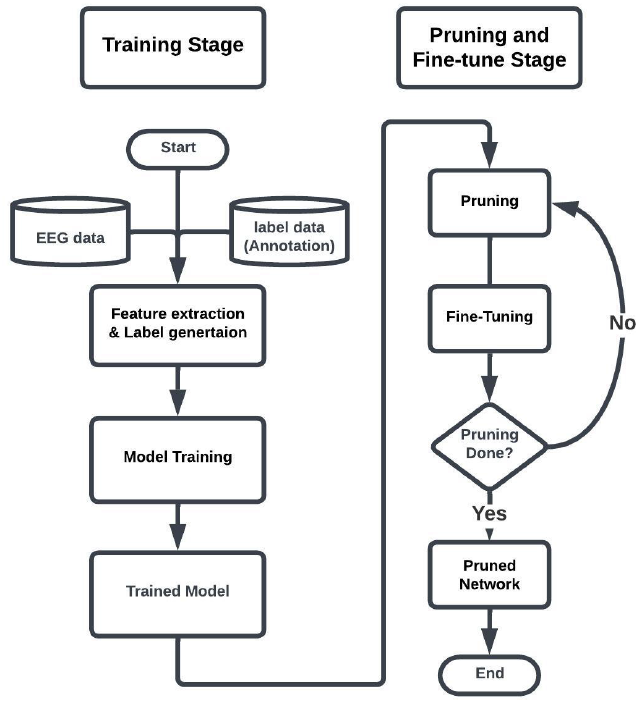
Flowchat of the training and pruning procedure; after the training completes, the pruning and fine-tuning are conducted iteratively to reduce the model size to save computational cost.

### D. Framework evaluation

#### 1) Evaluation patient cohort and iEEG recording

We evaluated the performance of the HFO detectors and classification by using the same data as in [10], [35]. Specifically, intracranial EEG (iEEG) data obtained via grid/strip electrodes using Nihon Kohden Systems (Neurofax 1100A, Irvine, California, USA) were used for analysis. . The study recording was acquired with a digital sampling frequency of 2,000 Hz. For each subject, separate 10-minute EEG segments from slow-wave sleep were selected at least two hours before or after seizures, before anti-seizure medication tapering, and before cortical stimulation mapping, which typically occurred two days after the implant. The annotation (Artifact, HFO-with-spike, HFO-without-spike) of each STE HFO event is also included in this data.

#### 2) HFO detector evaluation

To conduct a mathematical evaluation of the detection results between PyHFO and RIPPLELAB, we establish a defined representation of events detected by each algorithm. For PyHFO, an event is denoted as (*start*_*p*_, *end*_*p*_), indicating the exact time location within the EEG recording. Similarly, for RIPPLELAB, an event is represented as (*start*_*r*_, *end*_*r*_). To quantify the degree of overlap between these two sets of events, we introduce the concept of an overlapping ratio which is defined as 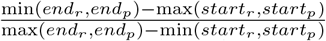. The resulting value ranges from 0 to 1, with 1 value indicating an exact match. To ensure a fair comparison and avoid double counting, we enforce the condition that an event detected by PyHFO can only match with a unique event detected by RIPPLELAB. Additionally, the comparison is performed on a channel-by-channel basis, and there is no overlap within events detected by the same detector by the definition of the detecting algorithm. The match of a specific event can be defined when the overlapping ratio exceeds a certain threshold, such as 50%. Furthermore, the discrepancy between the two algorithms can be quantified by calculating the ratio of the number of matches to the total number of events detected by RIPPLELAB.

We conducted four experiments to evaluate the success of our detector implementation, assessing the impact of each module independently in the pipeline. For simplicity, we denoted the data reading (Read), filter design (Filter), and detection algorithm (Algo) of RIPPLELAB as Read_*r*_ Filter_*r*_ and Algo_*r*_, where the subscript *r* represents the RIPPLELAB implementation, and we used the subscript *p* to denote the PyHFO implementation. To verify the correctness of our Python implementation, we first extracted the filtered EEG signal from RIPPLELAB. We fed it into our detectors (Read_*r*_ + Filter_*r*_ + Algo_*p*_), comparing the detection overlap with RIPPLELAB (Read_*r*_ + Filter_*r*_ + Algo_*r*_), which we refer to as Exp1. Since we replicated the logic of the two detectors, i.e., Algo_*r*_ = Algo_*p*_, we expected almost 100% matching between them. To assess the impact of the data reading, we conducted Exp2, replicating the frequency and phase response from RIPPLELAB in Python (Read_*p*_ + Filter_*r*_ + Algo_*p*_). For Exp3, we evaluated the effect of the filter design by feeding the EEG signal read by RIPPLELAB into our Python pipeline (Read_*r*_ + Filter_*p*_ + Algo_*p*_). Finally, we compared the complete implementation of PyHFO (Read_*p*_ + Filter_*p*_ + Algo_*p*_) with the RIPPLELAB implementation (Read_*r*_ + Filter_*r*_ + Algo_*r*_). We conducted all experiments on our 19 patients evaluation cohort and evaluated the implemented STE and MNI, respectively. For each experiment, we compared the detected HFOs with those detected from RIPPLELAB on the total number of HFOs, number of exact match HFOs, and the number of at least 50% overlap, evaluating the discrepancies between RIPPLELAB and our implementation in each step of the data processing. By evaluating the effect of each module independently, we were able to demonstrate the success of our detector implementation, providing a comprehensive assessment of the performance of our PyHFO implementation.

#### 3) DL-based neural network evaluation

The performance of our trained artifact and spk-HFO detectors was evaluated by comparing the results with expert labeling. For HFOs detected by the STE detector, we utilized the same dataset from [10], which consisted of 19 subjects and *n* = 12494 events. For HFOs identified by the MNI detector, two experts (NH and SH) randomly chose three subjects (*n* = 758), and annotated 416 artifacts, 312 spk-HFOs, and 30 non-spk-HFO. We adopted standard metrics for machine learning classification tasks to assess the model’s performance, including precision, recall, accuracy (Acc), and F1-score. Given that the model was trained using 5-fold cross-validation on STE HFOs, the reported metric values are the mean results of the 5-fold cross-validation (on different test sets) with a 95% confidence interval. For the MNI HFOs, models trained in 5-fold cross-validation were used to predict all events. The reported metric is thus the mean of the metrics from five models, again with a 95% confidence interval.

#### 4) Run time analysis

We conducted our run time analysis on different machines, Linux machine, Mac OS X machine and Windows machine. The Linux machine has an AMD Ryzen Threadripper 2950X 32-core processor; the Windows machine has an Intel i9-13900K 24-core; and the Mac OS X machine has an Apple M1 Pro 8-core processor. For timing the HFO detectors in RIPPLELAB, we follow the modification of RIPPLELAB in [10] to run the Matlab-based detector and report the run-time for detecting STE and MNI HFOs in the Linux machine. For PyHFO, we ran the detector with the same parameters as in RIPPLELAB and reported the run-time by using single-core (n-jobs =1) and multi-core (n-jobs = 32 for Linux, n-jobs = 8 for Windows and Mac machines). For benchmarking the DL models, we use DL models to predict 1000 samples ten times to get the mean and 95% confidence interval of the run time in different machines with PyTorch default setting, and we also report the inference time on an Nvidia Titan RTX GPU for reference.

### E. Software Overview

The software version PyHFO is a multi-window GUI developed in PyQt. It is intended to be a user-friendly and intuitive tool that users with technical and non-technical backgrounds can use to detect and classify HFOs in a time-efficient manner. PyHFO has been released under Academic Licenses (Licenses for Sharing Software Code Non-commercially, UCLA TDG). The source code and documentation of the PyHFO software application is open source in Github^1^. For technical background users; we also release our multi-processed based HFO detector in Python Package Index (PyPI) which can be installed by *pip install HFODetector* and DL-based HFO classifiers in Github ^2^ so that python users can easily install it. The GUI interface was implemented in PyQt version 5.15 to make it compatible with different hardware platforms such as OS X, Linux, and Windows. The HFO detectors are implemented in Python 3.9.0, and the DL-based detector is implemented in PyTorch 1.6. We chose Python as the programming platform because it is widely used in large-scale data analysis and deep learning in the medical image field. The completed procedure, for detecting and analyzing HFOs through PyHFO, consists of several steps briefly discussed in Figure 1. We here present the GUI of the software when the detection is done in Figure 7. After setting the parameters of different detectors and DL-based classifier, the original EEG signal will be displayed in the main visualization window, and HFO with different attributes (artifacts, HFO-with-spike, and HFO-without-spike) will be annotated with different colors. On the summary panel, the statistics of different kinds of HFOs will be displayed for each channel. All of these statistics can be exported in Excel format for further study by the user.

**Fig. 7.**
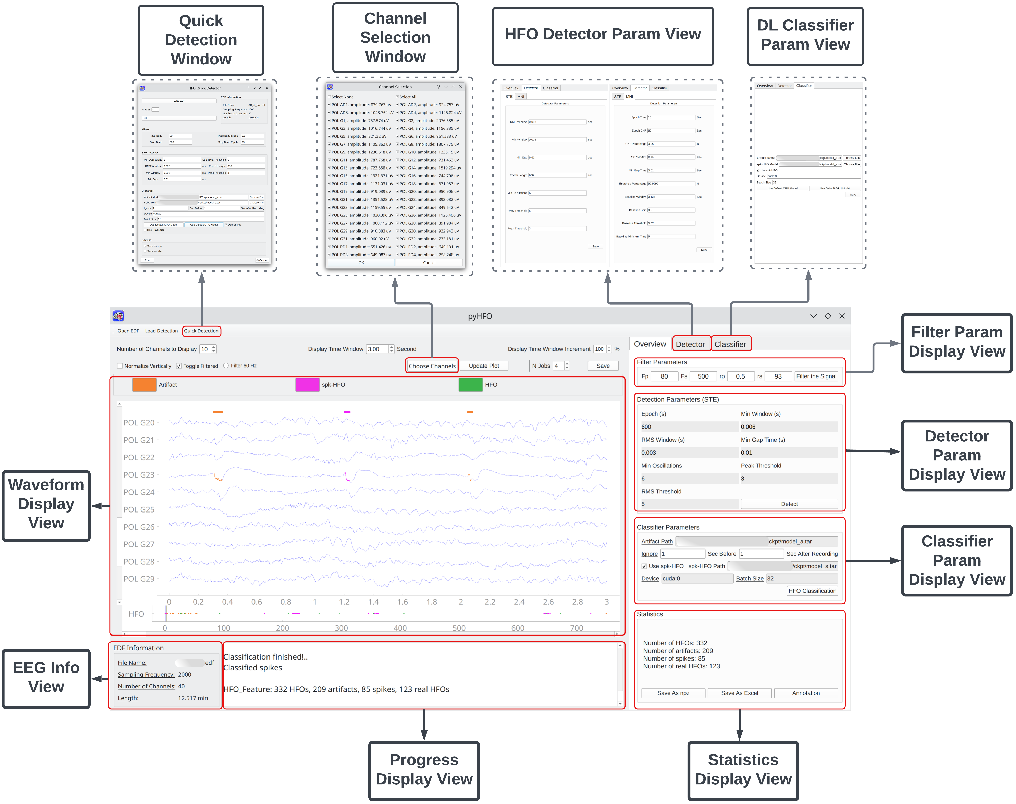
The multi-window overview of the PyHFO application, outlining the user interaction process. For a more detailed explanation, users should consult the PyHFO manual (reference needed). In summary, the process is as follows: **Load EDF**: The user can select and load an *.edf file for EEG data analysis. Basic EEG information and the EEG waveform are then displayed in the ‘EEG info view’ and ‘Waveform Display View’, respectively. **HFO detection**: Users can specify filter parameters in the ‘Filter Param Display View’, select the HFO detector and its parameters in the ‘HFO Detector Param View’, and click ‘detect’ to start HFO detection. The progress is displayed in the ‘Progress Display View’, and detected HFOs are shown in the ‘Waveform Display View’. **DL-based HFO classification**: Users can select a pre-trained network or use the pre-installed models in PyHFO from the ‘DL-Classifier Param View’. After clicking ‘HFO Classification’, a progress bar appears in the ‘Progress Display View’, and once the process completes, classified HFOs are marked in the ‘Waveform Display View’. Results can be exported in CSV or NPZ formats. To simplify the process, users can also use the ‘Quick Detection Window’ to specify all parameters for the whole pipeline, bypassing GUI interaction.

## III. Result

### A. HFO detector evaluation

Out of the 19 patients included in the study, a total of 1709 channels (with a median of 94 channels per patient) were analyzed using 10-minute EEG recordings from each patient. The RIPPLELAB by STE and MNI detectors detected 12, 494 and 10, 392 HFOs, while PyHFO detected 12, 501 and 10, 355, respectively. In Table I, we demonstrated the breakdown performance of each experiment. Specifically, in Exp1, when only the detection algorithm is different, we demonstrated that PyHFO successfully replicated the detection algorithms implemented in RIPPLELAB. The discrepancy in the MNI detector is due to different random seed RIPPLELAB and PyHFO used. Controlled variable experiments showed that different data readings (Exp2) and filter (Exp3) do affect the performance of the detector but with a minimal effect of around 3 to 7 % difference between RIPPLELAB’s detection and PyHFO. The overall discrepancy is defined as the sum of the number of new HFOs detected by the RIPPLELAB (new-RIPPLELAB) and the number of new HFOs detected by the PyHFO (new-PyHFO) divide the total number of HFOs detected by the RIPPLELAB. The discrepancies between PyHFO and RIPPLELAB of STE and MNI detector are 10% and 14%, respectively.

**TABLE I.** Comparison of Differences Between RippleLab and PyHFO Implementations.

|  | No. Events From STE Detector |  |  |  |  | No. Events From MNI Detector |  |  |  |  |
| --- | --- | --- | --- | --- | --- | --- | --- | --- | --- | --- |
|  | RIPPLELAB | Exp1 | Exp2 | Exp3 | PyHFO | RIPPLELAB | Exp1 | Exp2 | Exp3 | PyHFO |
| Total HFO | 12494 | 12494 | 12421 | 12582 | 12501 | 10392 | 10390 | 10320 | 10422 | 10355 |
| Exactly Same | - | 12494 | 11442 | 9147 | 8643 | - | 10368 | 10035 | 7729 | 7487 |
| 90% overlap | - | 12494 | 12192 | 12126 | 11876 | - | 10390 | 10136 | 9828 | 9612 |
| 50% overlap | - | 12494 | 12192 | 12144 | 11892 | - | 10390 | 10136 | 9835 | 9619 |
| new-RIPPLELAB | - | 0 | 302 | 350 | 602 | - | 2 | 256 | 557 | 773 |
| new-PyHFO | - | 0 | 229 | 438 | 609 | - | 0 | 184 | 587 | 736 |

As highlighted in Section II-B, the implementation within PyHFO closely adheres to prevailing methods for data reading. Additionally, it provides a more accurate representation of the input parameters utilized in the construction of the bandpass filter. Consequently, PyHFO’s methodology exhibits greater implementation accuracy compared to most mainstream publicly released software.

### B. HFO detector run-time comparison

Table II presents a runtime comparison between PyHFO and its Matlab-based counterpart across various hardware specifications. To save computational resources, we report only the runtime of RIPPLELAB on Linux machines. Based on the runtime comparison of PyHFO across different machines, it is evident that the runtime difference between RIPPLELAB’s operations on WindowsOS X and Linux is minimal.

**TABLE II.** Comparative Analysis of Runtime in RIPPLELAB and PyHFO: Detection of All Events from 10-Minute Data Segments Across 19 Patients (measured in minutes).

|  | Linux |  | Windows |  | OS X |  |
| --- | --- | --- | --- | --- | --- | --- |
|  | STE | MNI | STE | MNI | STE | MNI |
| RIPPLELAB | 372.83 | 5647.12 | - | - | - | - |
| PyHFO(n=1) | 57.43 | 971.35 | 34.57 | 933.31 | 35.90 | 659.63 |
| PyHFO(n>1) | 5.18 | 83.30 | 9.03 | 113.59 | 7.56 | 114.35 |

When comparing the runtime of HFO detection, PyHFO significantly outperforms RIPPLELAB in both single-core (n=1) and multi-core (n*>*1) configurations, as detailed in II-D4 for hardware setup specifications. The longest time required to detect all STE HFOs using PyHFO is only 9 minutes, compared to approximately 5 hours (373.83 minutes) when utilizing the best published HFO detector currently available.

Moreover, the longest time required for detecting MNI HFOs is 114 minutes with PyHFO, while RIPPLELAB takes approximately 4 days. Hence, PyHFO can be up to 50 times faster than mainstream Matlab-based software. Even when operating with a single core, PyHFO still offers at least a six-times improvement in speed.

Across various hardware configurations, the upper bound of the average time required to detect all STE HFOs for a single patient is around 30 seconds, and for the MNI detector, it is 17 minutes. The longer runtime for the MNI detector, compared to STE, is due to an iterative procedure within its computational pipeline. This enhanced performance allows for a comprehensive analysis of HFO in large patient groups and over extended recording periods.

### C. Machine learning algorithm against expert labeling

In five-fold cross-validation, for STE HFOs (19 subjects, n = 12494) the model achieved an accuracy of 98.6% and 89.1% for classifying artifacts and HFO with spikes, respectively, as shown in Table III. This performance is almost the same as that is [10] (artifacts: 98.8%, spk-HFO 89.1%) but with much lower MACs and run-time when we classify HFOs in CPUs in Table IV. More importantly, the model trained using STE HFOs can successfully classify the MNI HFOs (3 subjects, n = 758), demonstrating the success of the data augmentation and generalization-ability of the model, which enables these two deep learning models to be used in natural settings. The excellent performance across detectors also demonstrates the morphological similarity between the spk-HFO in MNI and STE detectors.

**TABLE III.** Performance Analysis using 5-Fold Cross-Validation: 95% Confidence Interval versus Expert Labeling.

|  | STE |  | MNI |  |
| --- | --- | --- | --- | --- |
|  | Artifacts | spk-HFO | Artifacts | spk-HFO |
| Accuracy | 98.56±0.19 | 89.18±0.82 | 98.72±0.64 | 89.79±3.12 |
| F1-score | 99.08±0.13 | 91.40±0.60 | 90.16±4.88 | 94.14±1.93 |
| Recall | 99.18±0.27 | 90.78±1.91 | 94.79±2.72 | 90.85±4.15 |
| Precision | 98.98±0.29 | 92.07±2.11 | 100.0±0.00 | 97.75±0.69 |

**TABLE IV.** Comparative Analysis of Computational Costs Across Models: MACs and Model Run-Time (in seconds) on GPU for Linux, OS X, and Windows.

|  | MACs | GPU | Linux | OS X | Windows |
| --- | --- | --- | --- | --- | --- |
| Artifact [10] | 1.82G | 0.14±0.10 | 9.78±0.20 | 20.28±1.89 | 15.68±0.10 |
| spk-HFO [10] | 1.82G | 0.14±0.11 | 9.84±0.25 | 19.43±1.16 | 16.01±0.74 |
| Artifact [ours] | 165.22M | 0.16±0.24 | 1.70±0.05 | 2.97±0.19 | 2.81±0.08 |
| spk-HFO[ours] | 312.90M | 0.07±0.01 | 2.70±0.11 | 4.13±0.20 | 3.98±0.05 |

### D. Neural network complexity comparison

In Table IV, we compare the Multiply Accumulate (MACs) on a single data sample as input and run-time of inference 1000 data samples using GPU and CPUs between state-of-the-art and PyHFO. We reported the performance metric of spk-HFO and artifact classifier, respectively. By computing the MACs, the classifiers in PyHFO are more computationally efficient than the models proposed in [10], which provide a theoretical support for later empirical experiments. Furthermore, even though both classifiers from [10] and PyHFO runs at comparable speed in GPU, the artifacts classifier in PyHFO runs at least 4 times faster than its counterpart, and PyHFO’s spk-HFO detector runs three times faster in CPUs. As another ablation study, we blindly pruned the published network using the same pruning and fine-tuning parameters but without input dimension optimization. Even though the performance of the pruned model is still comparable, MACs are still around 500M which is much higher than our approach.

## IV. Discussion

Our work was implemented based on the strong clinical motivation. Prior observational studies [35], [36] and a clinical trial [37] have shown issues with time constraints in HFO analysis in clinical settings. The clinical use of HFOs detection sites during epilepsy surgery planning requires a fast, reliable, and user-friendly application. It also needs to simulate human experts’ judgment to complement the entire process, including the detection and classification of HFOs. Our platform incorporated such capacities and has capacities to use multiple HFO detection methods and also has classifiers, including artifact rejection and HFOs with spikes vs. without spikes. Additionally, this system is portable, allowing any physician or researcher with a laptop to utilize DL-based algorithms in various settings, such as the epilepsy monitoring unit or the operating room. This capability has the potential to facilitate clinical trials. During the development of our PyHFO application, we demonstrated that our HFO detection algorithm is comparable with other open-source work, including RIPPLELAB. While we implemented the Python version of this HFO analysis application, EEG data reading and input format were deployed using the Python package. We followed the same EEG reading calibration as MNE [16], while in RIPPLELAB, the calibration was only done by voltage, without the offset adjustment. Furthermore, there were slight differences in the data filtering implementation. We choose to use the filter construction by Scipy as it can produce a more accurate frequency response. Our study reported minor differences in HFO detection numbers between RIPPLELAB and PyHFO, and we concluded that those are based on the aforementioned differences. We extend full credit to RIPPLELAB for developing the pioneering user-friendly, MATLAB-based HFO analysis software. This foundational effort greatly informed our Python-based platform and we also proved that our Python-based implementation is accurate based on engineering aspects.

We combined multiple methods to decrease the run time of the whole pipeline. We utilized the multi-processing feature of Python, employed vectorization implementation in wavelet computation, optimized the neural network’s input size and pruned the neural network architecture. We also developed a data-augmentation strategy in the neural network training to improve the generalization ability of the model. We demonstrated that with the use of our application, the run-time was about 50 times faster in STE detection and MNI detection compared to RIPPLELAB. We also achieved high performance in classifying artifacts and HFOs with spikes (98.6% and 89.1%, respectively).

There are several limitations to our study. Our application was developed using iEEG data sampled from grids/strips, not from SEEG. Additionally, our subjects are pediatric patients with various epilepsy etiologies. Moreover, our data is from a single institution. The generalizability of our application is still considered limited as it stands. However, our application has the potential to expand its capacity.

In the near future, we have plans to incorporate a diverse dataset into our system. Our objective is to test the system on a larger dataset comprising over 100 subjects, including pediatric and adult patient data acquired through grids/strips and SEEG. The versatility of our application is evident as we strive to incorporate additional detection methods, such as Hilbert [38] or SLL [39]. As we continuously expand the dataset and introduce new functionalities, the algorithm’s performance will progressively improve through training. The invaluable real-time feedback from frontline physicians and researchers will contribute significantly to this iterative process.

Moreover, our integration of the DL-based algorithm aims to have a broader scientific impact. This integration empowers us to undertake more challenging tasks, such as distinguishing between pathological and physiological HFOs, and leveraging DL to characterize distinctive features for each HFO type. The potential of our platform extends further, enabling seamless collaboration and facilitating large-scale EEG data analysis. This breakthrough opens up new avenues for scientific exploration in the field.

## V. ACKNOWLEDGMENT

The authors have no conflict of interest to disclose. HN is supported by the National Institute of Neurological Disorders and Stroke (NINDS) K23NS128318, the Sudha Neelakantan & Venky Harinarayan Charitable Fund, the Elsie and Isaac Fogelman Endowment, and the UCLA Children’s Discovery and Innovation Institute (CDI) Junior Faculty Career Development Grant (#CDI-TTCF-07012021). AD is supported to research abroad by the Uehara Memorial Foundation and SENSHIN Medical Research Foundation. SAH has received research support from the Epilepsy Therapy Project, the Milken Family Foundation, the Hughes Family Foundation, the Elsie and Isaac Fogelman Endowment, Eisai, Lundbeck, Insys, Zogenix, GW Pharmaceuticals, UCB, and has received honoraria for service on the scientific advisory boards of Questcor, Mallinckrodt, Insys, UCB, and Upsher-Smith, for service as a consultant to Eisai, UCB, GW Pharmaceuticals, Insys, and Mallinckrodt, and for service on the speakers’ bureaus of Mallinckrodt and Greenwich Bioscience. RS serves on scientific advisory boards and speakers bureaus and has received honoraria and funding for travel from Eisai, Greenwich Biosciences, UCB Pharma, Sunovion, Supernus, Lundbeck Pharma, Liva Nova, and West Therapeutics (advisory only); receives royalties from the publication of Pellock’s Pediatric Neurology (Demos Publishing,2016) and Epilepsy: Mechanisms, Models, and Translational Perspectives (CRC Press, 2011). RJS is supported by the National Institute of Neurological Disorders and Stroke (NINDS) R01NS106957 and Christina Louise George Trust.

## Appendix A

### Details of two detection algorithms

#### 1) Short Time Energy HFO detection algorithm

The first implemented algorithm, Short Time Energy (STE), detects HFO events by selecting the energy of the filtered raw EEG signal with an estimated energy threshold for each 10-min epoch. In detail, the EEG signal is processed through a bandpass filter in frequencies between 80 and 500 Hz. The energy of the filtered signal is then computed based on RMS with N = 3 ms window defined by the equation

The estimation of the energy threshold is 5 standard deviations (SD) above the overall RMS mean. Finally, all HFO events selected should be with a duration of more than 6 ms and contain more than 6 peaks greater than 3 SD above the filtered signal mean value.

#### 2) Montreal Neurological Institute’s HFO detection algorithm

Another implemented algorithm MNI detector (MNI) was proposed by Zelmann et al. For this approach, similar to the STE algorithm, the raw EEG signal is also filtered by the bandpass filter and the energy of the filtered signal is then computed using root mean square (RMS) with 2 ms window. A key block for MNI algorithm is the baseline detector, designed to construct the baseline interval. The baseline, defined as EEG segments with no oscillation, is detected by computing the wavelet entropy of the autocorrelation of the filtered signal. For each 125 ms EEG segment with 50% shift, the segment is considered as baseline if the wavelet entropy is greater than the threshold. From the baseline detector, possible HFO events could be detected by selecting energy of the filtered signal using RMS above the energy threshold for each epoch. The energy threshold is computed in two different methods if more than or less than 5 s/min of the amount of all detected baseline. If sufficient baseline was found, the energy threshold is selected from RMS values for each 10s baseline segment at the 99.9999 percentile of its empirical cumulative distribution function. If less baseline was detected, we consider the channel with continuous high-frequency oscillatory activity. The energy threshold is iteratively selected from RMS values for each 60s segment at the 95 percentile of its empirical cumulative distribution function. The value is found by continuously detecting and removing the highest energy till no more new HFO events are detected. All possible HFO events from the two situations are finally selected with a duration of more than 6 ms.

## Footnotes

1 https://github.com/roychowdhuryresearch/pyHFO

2 https://github.com/roychowdhuryresearch/HFO-Classification/tree/main/Pruning-pipeline

## References

[1] A. Bragin, J. Engel Jr, C. L. Wilson, I. Fried, and G. W. Mathern, “Hippocampal and entorhinal cortex high-frequency oscillations (100– 500 hz) in human epileptic brain and in kainic acid-treated rats with chronic seizures,” Epilepsia, vol. 40, no. 2, pp. 127–137, 1999.

[2] R. J. Staba, C. L. Wilson, A. Bragin, D. Jhung, I. Fried, and J. Engel Jr, “High-frequency oscillations recorded in human medial temporal lobe during sleep,” Annals of neurology, vol. 56, no. 1, pp. 108–115, 2004.

[3] J. Jirsch, E. Urrestarazu, P. LeVan, A. Olivier, F. Dubeau, and J. Gotman, “High-frequency oscillations during human focal seizures,” Brain, vol. 129, no. 6, pp. 1593–1608, 2006.

[4] G. A. Worrell, L. Parish, S. D. Cranstoun, R. Jonas, G. Baltuch, and B. Litt, “High-frequency oscillations and seizure generation in neocortical epilepsy,” Brain, vol. 127, no. 7, pp. 1496–1506, 2004.

[5] T. Akiyama, B. McCoy, C. Y. Go, A. Ochi, I. M. Elliott, M. Akiyama, J. Donner, S. K. Weiss, O. C. Snead III, J. T. Rutka et al., “Focal resection of fast ripples on extraoperative intracranial eeg improves seizure outcome in pediatric epilepsy,” Epilepsia, vol. 52, no. 10, pp. 1802–1811, 2011.

[6] J. Wu, R. Sankar, J. Lerner, J. Matsumoto, H. Vinters, and G. Mathern, “Removing interictal fast ripples on electrocorticography linked with seizure freedom in children,” Neurology, vol. 75, no. 19, pp. 1686–1694, 2010.

[7] J. Jacobs, M. Zijlmans, R. Zelmann, C.-É. Chatillon, J. Hall, A. Olivier Dubeau, and J. Gotman, “High-frequency electroencephalographic oscillations correlate with outcome of epilepsy surgery,” Annals of Neurology: Official Journal of the American Neurological Association and the Child Neurology Society, vol. 67, no. 2, pp. 209–220, 2010.

[8] M. A. van’t Klooster, N. E. van Klink, W. J. Zweiphenning, F. S. Leijten, R. Zelmann, C. H. Ferrier, P. C. van Rijen, W. M. Otte, K. P. Braun, G. J. Huiskamp et al., “Tailoring epilepsy surgery with fast ripples in the intraoperative electrocorticogram,” Annals of neurology, vol. 81, no. 5, pp. 664–676, 2017.

[9] T. Monsoor, Y. Zhang, A. Daida, S. Oana, Q. Lu, S. A. Hussain, A. Fallah, R. Sankar, R. J. Staba, W. Speier, V. Roychowdhury, and H. Nariai, “Optimizing detection and deep learning-based classification of pathological high-frequency oscillations in epilepsy,” Clinical Neurophysiology, 2023. [Online]. Available: https://www.sciencedirect.com/science/article/pii/S1388245723006971

[10] Y. Zhang, Q. Lu, T. Monsoor, S. A. Hussain, J. X. Qiao, N. Salamon, A. Fallah, M. S. Sim, E. Asano, R. Sankar et al., “Refining epileptogenic high-frequency oscillations using deep learning: a reverse engineering approach,” Brain communications, vol. 4, no. 1, p. fcab267, 2022.

[11] Y. Zhang, H. Chung, J. P. Ngo, T. Monsoor, S. A. Hussain, J. H. Matsumoto, P. D. Walshaw, A. Fallah, M. S. Sim, E. Asano et al., “Characterizing physiological high-frequency oscillations using deep learning,” Journal of neural engineering, vol. 19, no. 6, p. 066027, 2022.

[12] A. Delorme and S. Makeig, “Eeglab: an open source toolbox for analysis of single-trial eeg dynamics including independent component analysis,” Journal of neuroscience methods, vol. 134, no. 1, pp. 9–21, 2004.

[13] M. Hassan, M. Shamas, M. Khalil, W. El Falou, and F. Wendling, “Eegnet: An open source tool for analyzing and visualizing m/eeg connectome,” PloS one, vol. 10, no. 9, p. e0138297, 2015.

[14] D. Currey, J. Craley, D. Hsu, R. Ahmed, and A. Venkataraman, “Epviz: A flexible and lightweight visualizer to facilitate predictive modeling for multi-channel eeg,” Plos one, vol. 18, no. 2, p. e0282268, 2023.

[15] S. Baillet, K. Friston, and R. Oostenveld, “Academic software applications for electromagnetic brain mapping using meg and eeg,” Computational intelligence and neuroscience, vol. 2011, pp. 12–12, 2011.

[16] A. Gramfort, M. Luessi, E. Larson, D. A. Engemann, D. Strohmeier, C. Brodbeck, R. Goj, M. Jas, T. Brooks, L. Parkkonen, and M. S. Hämäläinen, “MEG and EEG data analysis with MNE-Python,” Frontiers in Neuroscience, vol. 7, no. 267, pp. 1–13, 2013.

[17] R. Vallat and M. P. Walker, “An open-source, high-performance tool for automated sleep staging,” Elife, vol. 10, p. e70092, 2021.

[18] F. S. Bao, X. Liu, C. Zhang et al., “Pyeeg: an open source python module for eeg/meg feature extraction,” Computational intelligence and neuroscience, vol. 2011, 2011.

[19] D. Jacobs, Y. Liu, T. Hilton, M. Campo, P. Carlen, and B. Bardakjian, “Classification of scalp eeg states prior to clinical seizure onset,” IEEE Journal of Translational Engineering in Health and Medicine, vol. PP, pp. 1–1, 08 2019.

[20] S. Wang, I. Z. Wang, J. C. Bulacio, J. C. Mosher, J. Gonzalez-Martinez, A. V. Alexopoulos, I. M. Najm, and N. K. So, “Ripple classification helps to localize the seizure-onset zone in neocortical epilepsy,” Epilepsia, vol. 54, no. 2, pp. 370–376, 2013. [Online]. Available: https://onlinelibrary.wiley.com/doi/abs/10.1111/j.1528-1167.2012.03721.x

[21] W. Yin, Z. Liang, J. Zhang, and Q. Liu, “Partial least square regression via three-factor svd-type manifold optimization for eeg decoding,” in Chinese Conference on Pattern Recognition and Computer Vision (PRCV). Springer, 2022, pp. 778–787.

[22] H. Zhang, M. Zhao, C. Wei, D. Mantini, Z. Li, and Q. Liu, “Eegdenoisenet: a benchmark dataset for deep learning solutions of eeg denoising,” Journal of Neural Engineering, vol. 18, no. 5, p. 056057, 2021.

[23] M. Lu, Y. Zhang, A. Daida, S. Oana, R. R. Rajaraman, H. Nariai, and S. A. Hussain, “Application of an eeg-based deep learning model to discriminate children with epileptic spasms from normal controls,” medRxiv, pp. 2023–06, 2023.

[24] M. Navarrete, C. Alvarado-Rojas, M. Le Van Quyen, and M. Valderrama, “Ripplelab: A comprehensive application for the detection, analysis and classification of high frequency oscillations in electroencephalographic signals,” PloS one, vol. 11, no. 6, p. e0158276, 2016.

[25] H. Nariai, S. A. Hussain, D. Bernardo, H. Motoi, M. Sonoda, N. Kuroda, E. Asano, J. C. Nguyen, D. Elashoff, R. Sankar et al., “Scalp eeg interictal high frequency oscillations as an objective biomarker of infantile spasms,” Clinical Neurophysiology, vol. 131, no. 11, pp. 2527–2536, 2020.

[26] S. V. Gliske, Z. T. Irwin, C. Chestek, G. L. Hegeman, B. Brinkmann, O. Sagher, H. J. Garton, G. A. Worrell, and W. C. Stacey, “Variability in the location of high frequency oscillations during prolonged intracranial eeg recordings,” Nature communications, vol. 9, no. 1, p. 2155, 2018.

[27] N. Kuroda, M. Sonoda, M. Miyakoshi, H. Nariai, J.-W. Jeong, H. Motoi, A. F. Luat, S. Sood, and E. Asano, “Objective interictal electrophysiology biomarkers optimize prediction of epilepsy surgery outcome,” Brain communications, vol. 3, no. 2, p. fcab042, 2021.

[28] C. P. Lisgaras and H. E. Scharfman, “High-frequency oscillations (250– 500 hz) in animal models of alzheimer’s disease and two animal models of epilepsy,” Epilepsia, vol. 64, no. 1, pp. 231–246, 2023.

[29] K. J. Barth, J. Sun, C.-H. Chiang, S. Qiao, C. Wang, S. Rahimpour, M. Trumpis, S. Duraivel, A. Dubey, K. E. Wingel et al., “Flexible, high-resolution cortical arrays with large coverage capture microscale high-frequency oscillations in patients with epilepsy,” Epilepsia, 2023.

[30] G. T. Petito, J. Housekeeper, J. Buroker, C. Scholle, B. Ervin, C. Frink, H. M. Greiner, J. Skoch, F. T. Mangano, T. J. Dye et al., “Diurnal rhythms of spontaneous intracranial high-frequency oscillations,” Seizure, vol. 102, pp. 105–112, 2022.

[31] A. Navas-Olive, R. Amaducci, M.-T. Jurado-Parras, E. R. Sebastian, and L. M. de la Prida, “Deep learning-based feature extraction for prediction and interpretation of sharp-wave ripples in the rodent hippocampus,” Elife, vol. 11, p. e77772, 2022.

[32] R. J. Staba, C. L. Wilson, A. Bragin, I. Fried, and J. Engel Jr, “Quantitative analysis of high-frequency oscillations (80–500 hz) recorded in human epileptic hippocampus and entorhinal cortex,” Journal of neurophysiology, vol. 88, no. 4, pp. 1743–1752, 2002.

[33] R. Zelmann, F. Mari, J. Jacobs, M. Zijlmans, R. Chander, and J. Gotman, “Automatic detector of high frequency oscillations for human recordings with macroelectrodes,” in 2010 Annual International Conference of the IEEE Engineering in Medicine and Biology. IEEE, 2010, pp. 2329–2333.

[34] D. Traxl, N. Boers, and J. Kurths, “Deep graphs—A general framework to represent and analyze heterogeneous complex systems across scales,” Chaos: An Interdisciplinary Journal of Nonlinear Science, vol. 26, no. 6, p. 065303, 06 2016. [Online]. Available: 10.1063/1.4952963

[35] H. Nariai, S. A. Hussain, D. Bernardo, A. Fallah, K. K. Murata, J. C. Nguyen, R. R. Rajaraman, L. M. Rao, J. H. Matsumoto, J. T. Lerner et al., “Prospective observational study: Fast ripple localization delineates the epileptogenic zone,” Clinical neurophysiology, vol. 130, no. 11, pp. 2144–2152, 2019.

[36] J. Jacobs, J. Y. Wu, P. Perucca, R. Zelmann, M. Mader, F. Dubeau, G. W. Mathern, A. Schulze-Bonhage, and J. Gotman, “Removing high-frequency oscillations: A prospective multicenter study on seizure outcome,” Neurology, vol. 91, no. 11, pp. e1040–e1052, 2018.

[37] W. Zweiphenning, M. A. van’t Klooster, N. E. van Klink, F. S. Leijten, C. H. Ferrier, T. Gebbink, G. Huiskamp, M. J. van Zandvoort, M. M. van Schooneveld, M. Bourez et al., “Intraoperative electrocorticography using high-frequency oscillations or spikes to tailor epilepsy surgery in the netherlands (the hfo trial): a randomised, single-blind, adaptive non-inferiority trial,” The Lancet Neurology, vol. 21, no. 11, pp. 982–993, 2022.

[38] B. Crépon, V. Navarro, D. Hasboun, S. Clemenceau, J. Martinerie, M. Baulac, C. Adam, and M. Le Van Quyen, “Mapping interictal oscillations greater than 200 hz recorded with intracranial macroelectrodes in human epilepsy,” Brain, vol. 133, no. 1, pp. 33–45, 2010.

[39] A. B. Gardner, G. A. Worrell, E. Marsh, D. Dlugos, and B. Litt, “Human and automated detection of high-frequency oscillations in clinical intracranial eeg recordings,” Clinical neurophysiology, vol. 118, no. 5, pp. 1134–1143, 2007.

